# Rapid and real-time identification of fungi up to the species level with long amplicon Nanopore sequencing from clinical samples

**DOI:** 10.1101/2020.02.06.936708

**Authors:** Sara D’Andreano, Anna Cuscó, Olga Francino

## Abstract

The availability of long-read technologies, like Oxford Nanopore Technologies, provides the opportunity to sequence longer fragments of the fungal ribosomal operon, up to 6 Kb (18S-ITS1-5.8S-ITS2-28S), and to improve the taxonomy assignment of the communities up to the species level and in real-time. We assess the taxonomy skills of amplicons targeting a 3.5 Kb region (V3 18S-ITS1-5.8S-ITS2-28S D2) and a 6 Kb region (V1 18S-ITS1-5.8S-ITS2-28S D12) with the What’s in my pot (WIMP) classifier. We used the ZymoBIOMICS™ mock community and different microbiological fungal cultures as positive controls. Long amplicon sequencing correctly identified *Saccharomyces cerevisiae* and *Cryptococcus neoformans* from the mock community and *Malassezia pachydermatis, Microsporum canis*, and *Aspergillus fumigatus* from the microbiological cultures. Besides, we identified *Rhodotorula graminis* in a culture mislabeled as *Candida* spp.

We applied the same approach to external otitis in dogs. *Malassezia* was the dominant fungal genus in dogs’ ear skin, whereas *M. pachydermatis* was the main species in the healthy sample. Conversely, we identified a higher representation of *M. globosa* and *M. sympodialis* in otitis affected samples. We demonstrate the suitability of long ribosomal amplicons to characterize the fungal community of complex samples, either healthy or with clinical signs of infection.

## INTRODUCTION

Fungi is a vast kingdom of organisms with a range between 1.5 and 6 million species (Hawksworth and Lücking 2017), but only a modest part, around 140.000 species, is phenotypical and genetically described (Hibbett et al. 2016; Wurzbacher et al. 2018). Usually, fungi have been identified by morphology or pure cultures in agar medium. The main problem is that many species are difficult to isolate and culture, and even to classify (Arbefeville et al. 2017; Usyk et al. 2017).

Both clinical approaches and research applications need taxonomic classification to assign taxa to their functional traits (Dayarathne et al. 2016; Raja et al. 2017). On the other hand, problems of unknown branches of fungal phylogenies still occur, due to considerable gaps in genetic knowledge and to old species description (Tedersoo et al. 2018).

Sequence-based methods allow to classify the fungi kingdom better. Still, the choice of the methodology used to study the mycobiome, or even the intrinsic characteristics of a specific fungus, can impact the data generated and the results reached (Usyk et al. 2017). Public online databases for fungal identification are noteworthy but still limited due to new updates for the best nomenclature and identification of fungi species (Prakash et al. 2017). A large number of online fungal databases are available for mycology research, and they are growing thanks to the dedication of the experts’ team. Taxonomic revisions are still ongoing and the main databases for fungi classification are Species Fungorum (www.speciesfungorum.org), MycoBank (www.mycobank.org), UNITE (Abarenkov et al. 2010) and International Nucleotide Sequence Database Consortium (https://www.ncbi.nlm.nih.gov/taxonomy).

One of the preferred markers for taxonomy assignment is the fungal ribosomal operon, which is almost 6.000 bp length. It contains three conserved units, the 18S rRNA gene (small subunit, SSU), 5.8S rRNA gene, and 28S rRNA gene (large subunit, LSU), and two hypervariable units as internal transcribed spacers regions (ITS1 and ITS2). The ITS units flank the 5.8S RNA gene, and better represent the high variability among taxonomic levels of fungi; showing a superior species discrimination and PCR success rates (Kalan and Grice 2017). The variable domains located at the conserved 18S (V1-V9) and 28S rRNA genes (D1-D12) (Figure 1) are also worth considering to refine the taxonomy assignment. It is essential to recognize that the D1-D2 from the 28S rRNA gene domains are the ones that perform a higher level of taxonomic assignment for fungi (Tedersoo et al. 2015).

**Figure 1.**
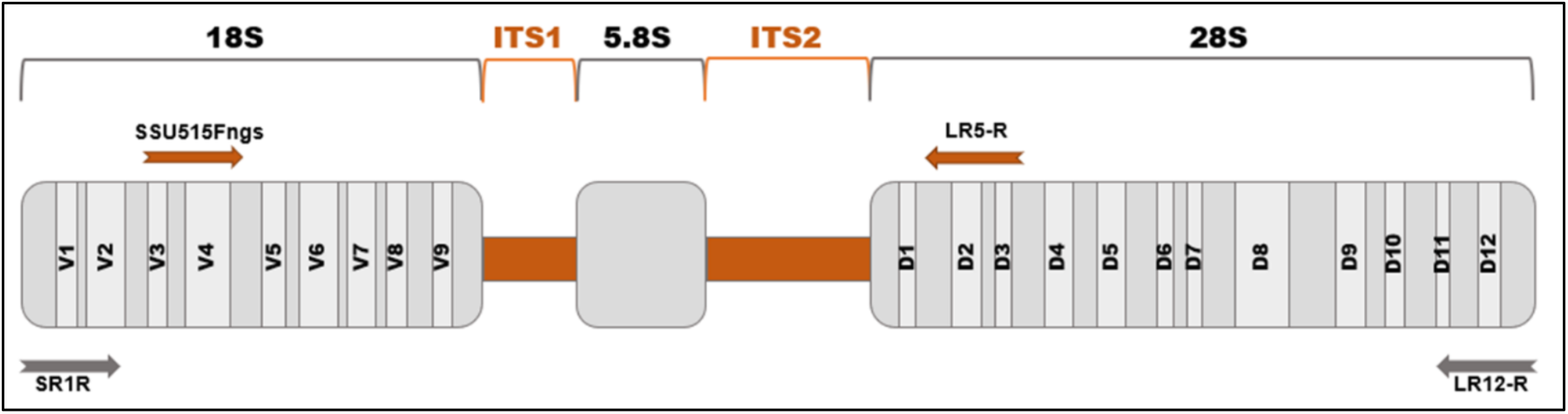
Fungal ribosomal operon: two hypervariable internal transcribed spacers regions (ITS1 and ITS2, marked in orange) and three conserved ones (18S, 5.8S, and 28S rRNA, marked in grey) that contain variables domains, nine for the 18S and twelve for the 28S rRNA genes. Primers set used for the amplification of the ITS region (3.5 Kb) are shown in orange in the upper part of the operon, and the ones for amplification of the full operon (6 Kb) in grey in the lower part (Vilgalys lab 1992; Tedersoo et al. 2015)

Primer sets to amplify the fungal operon regions have been described in different manuscripts, starting from 1990 until 2018 (White et al. 1990; Vilgalys lab 1992; Lee et al. 2008; Ihrmark et al. 2012; Tedersoo and Lindahl 2016; Tedersoo et al. 2018). These sets of primers target the appropriate operon fragments to proceed with either short fragments, by massively parallel sequencing (or 2^nd^ generation sequencing) as Ion Torrent or Illumina, or longer fragments with single-molecule sequencing (or 3^rd^ generation sequencing) as PacBIO or Oxford Nanopore Technologies.

Taxonomy with short reads is focused on ITS1 and ITS2 regions, considered as the official barcoding markers for species-level identification in fungi, due to their easy amplification, conserved primers sites, widespread use (Schoch et al. 2012) and available databases, such as UNITE or Mycobank. Usually, the ITS1 and ITS2 regions provide the taxonomy resolution within-genus and within-species level, but debates on which one provides the best taxonomic skill are still under discussion (Blaalid et al. 2013). Alternative markers located in the small and large subunits of the rRNA genes can address the phylogenetic diversity, depending on the fungi species (Blaalid et al. 2013; Tedersoo et al. 2015). Depending on the fungus species, different regions of the operon can be considered for taxonomy assignment: the SSU and the LSU are used when taxonomy is investigated up to family level, while lower taxonomy level analysis requires the ITS regions. When primers sets include the D1-D2 regions of LSU subunit, fragments obtained from the amplification can be assigned up to the species level (Raja et al. 2017).

Here, we aim to investigate the taxonomic skills of the long-amplicon PCR approach to detect and identify the fungal microbiota present on complex microenvironments at the species level. We use microbiological fungal cultures characterized phenotypically as positive controls. We then apply the same protocol to clinical samples of canine otitis as a complex microenvironment.

## MATERIALS and METHODS

### Samples and DNA extraction

LETI laboratories (LETI Animal Health) kindly provided a total of eight microbiological fungal cultures in Petri dishes. Four of the cultures had been classified up to the genus level (*Alternaria* spp, *Aspergillus* spp, *Candida* spp and *Malassezia* spp) and four other ones up to the species level: three of *Malassezia pachydermatis* and one of *Microsporum canis.* Also, fungal DNA of the ZymoBIOMICS™ mock community containing *Saccharomyces cerevisiae* and *Cryptococcus neoformans* was included in the study as a positive control. The DNA from all fungal samples was extracted by ZymoBIOMICS™ Miniprep kit following the manufacturer’s instructions.

Four canine otitis samples were analysed as complex microbial microenvironments. Two of them were collected from a Petri dish, divided into two halves parts, to culture fungi from both ears of a dog, one of the ears was healthy, and the other one showed clinical signs for external otitis (S02_healthy; S03_affected). The DNA was extracted by using ZymoBIOMICS™ Miniprep kit, as for the cultures. The other two otitis samples (S01_affected; S04_affected) were collected by swabbing the inner pinna of the ear of two dogs using Sterile Catch-All™ Sample Collection Swabs (Epicentre Biotechnologies). DNA was extracted with QIAGEN-DNeasy PowerSoil Kit. DNA quality control was checked by Nanodrop and Qubit™ Fluorometer (Life Technologies, Carlsbad, CA).

### MinION sequencing

Two sets of primers were chosen (Table 1; Figure 1): the first set targeting the ribosomal operon from V3 region of 18S RNA gene to D3 region of 28S RNA gene (≈3.500 bp), and the second one targeting the complete ribosomal operon from V1 region of 18S RNA gene to D12 region of 28S RNA gene (≈6000 bp). The primers, both forward and reverse, included the Nanopore Universal Tags.

**Table 1.**
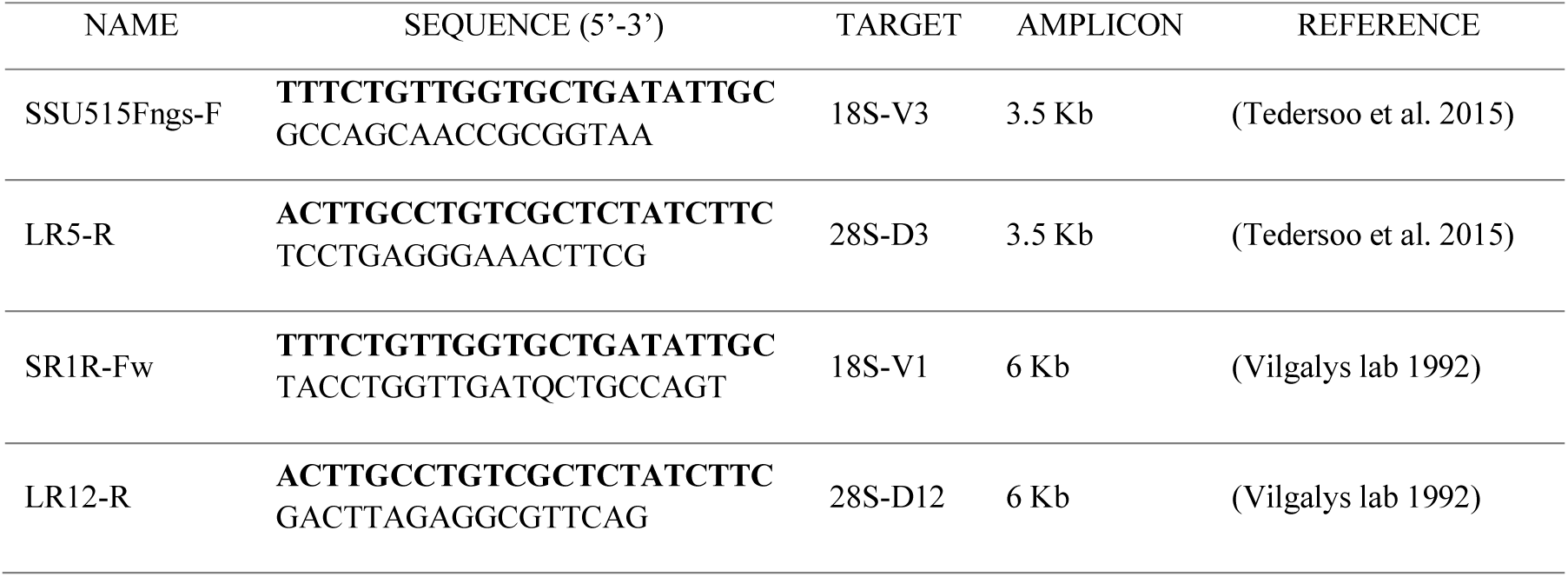
Primers targeting the full ITS region (3.5 Kb) and the whole fungal operon (6 Kb). The Nanopore Universal Tag in bold type.

Two PCRs were performed: the first for the amplification of the target, and the second one to add the specific barcode to each sample. DNA initial concentration was of 5 ng DNA per sample, in 50 µl of PCR final volume: 15 µl of DNA plus 35 µl of PCR mix, which contained 10 µl of Phusion® High Fidelity Buffer (5x), 5 µl of dNTPs (2 mM), 0.5 µM of primer forward and reverse, and 0.02 U/µl of Phusion® Hot Start II Taq Polymerase (Thermo Scientific). PCR profile included an initial denaturation of 30 s at 98 °C, followed by 25 cycles of 10 s at 98 °C, 30 s at 62 °C, 80 s at 72 °C, and a final extension of 10 min at 72 °C. Amplicons obtained were purified with Agencourt AMPure XP beads, at 0.4X ratio for the fungal amplicon; then, they were quantified by Qubit™ fluorometer (Life Technologies, Carlsbad, CA).

Following the PCR Barcoding kit protocol (SQK-PBK004), 0.5 nM per each sample was required for the second PCR to add barcodes of PCR barcoding kit (EXP-PBC001). S The final volume of second PCR is 100 µl, containing 20 µl of DNA template from the previous PCR at 0.5 nM, 2 µl of specific barcode and 78 µl of mixture that include: 20 µl of 5× Phusion® High Fidelity Buffer, 10 µl of dNTPs (2 mM) and 2 U/µl of Phusion® Hot Start II Taq Polymerase. PCR profile included an initial denaturation of 30 s at 98 °C, 15 cycles of 10 s at 98 °C, 30 s at 62 °C, 80 s at 72 °C and final step of 10 min at 72 °C. The amplicon obtained were purified again with Agencourt AMPure XP beads, at 0.4X ratio and quantified by Qubit™ fluorometer (Life Technologies, Carlsbad, CA).

We proceeded then to the Library preparation with the Ligation Sequencing kit (SQK-LSK109, Oxford Nanopore Technologies). Barcoded samples (1.5 µg) were pooled in 47 µl of nuclease-free water and the library was prepared following the manufacturer conditions.

With a final step of Agencourt AMPure XP beads 0.4X, the DNA library was cleaned and ready to be loaded into the flow cell. We used two SpotON Flow Cells (FLO-MIN106) for three MinION runs, primed with a mixture of sequencing buffer and Flush buffer according to the manufacturer’s instructions. A quality control of sequencing pores was done before each run. Libraries were mixed with Sequencing Buffer and Loading Beads in a final volume of 75 µl. The final mix was added, by dropping, in the SpotON sample port.

Sequencing runs were between 16h and 19h, using the MinKNOWN 2.2 v18.07.2 and the MinKNOWN v18.12.9.

### Data Analysis

The sequencing of the 3.5 Kb amplicon generated fast5 files that were basecalled and demultiplexed by Albacore v2.3.3 for the 3.5 Kb amplicon or guppy 2.3.5 for the 6 Kb amplicon. Using Porechop (https://github.com/rrwick/Porechop), barcodes and adapters were removed. For the taxonomy assignment, we applied the cloud-based analysis *What’s in my pot* (WIMP) application from the EPI2ME platform (Methricor), based in Centrifuge (https://ccb.jhu.edu/software/centrifuge/manual.shtml).

The fastq files output of each fungal amplicon with the length of 3.5 Kb and 6 Kb, are loaded on Zenodo (http://doi.org/10.5281/zenodo.3662300).

## RESULTS

We aim to develop a long-amplicon PCR approach to detect and identify fungal microbiota present on complex microenvironments, and to apply it to clinical samples (canine otitis). As positive controls, we chose microbiological fungal cultures and fungal strains from a mock community. Some of the cultures were previously classified by classical microbiology up to the genus level as *Alternaria* spp, *Aspergillus* spp, *Candida* spp, and *Malassezia* spp. Others were classified up to the species level as *Malassezia pachydermatis* and *Microsporum canis* at LETI laboratories (LETI Animal Health). The ZymoBIOMICS™ mock community fungal strains are *Saccharomyces cerevisiae* and *Cryptococcus neoformans.*

### Identification of microbiological cultures and mock community with long amplicons

All samples were amplified for both amplicon sizes, 3.5 Kb and 6 Kb. In the 3.5 Kb amplicons, we included those domains that better help in the taxonomic classification of fungi, as shown in Figure 2.

**Figure 2:**
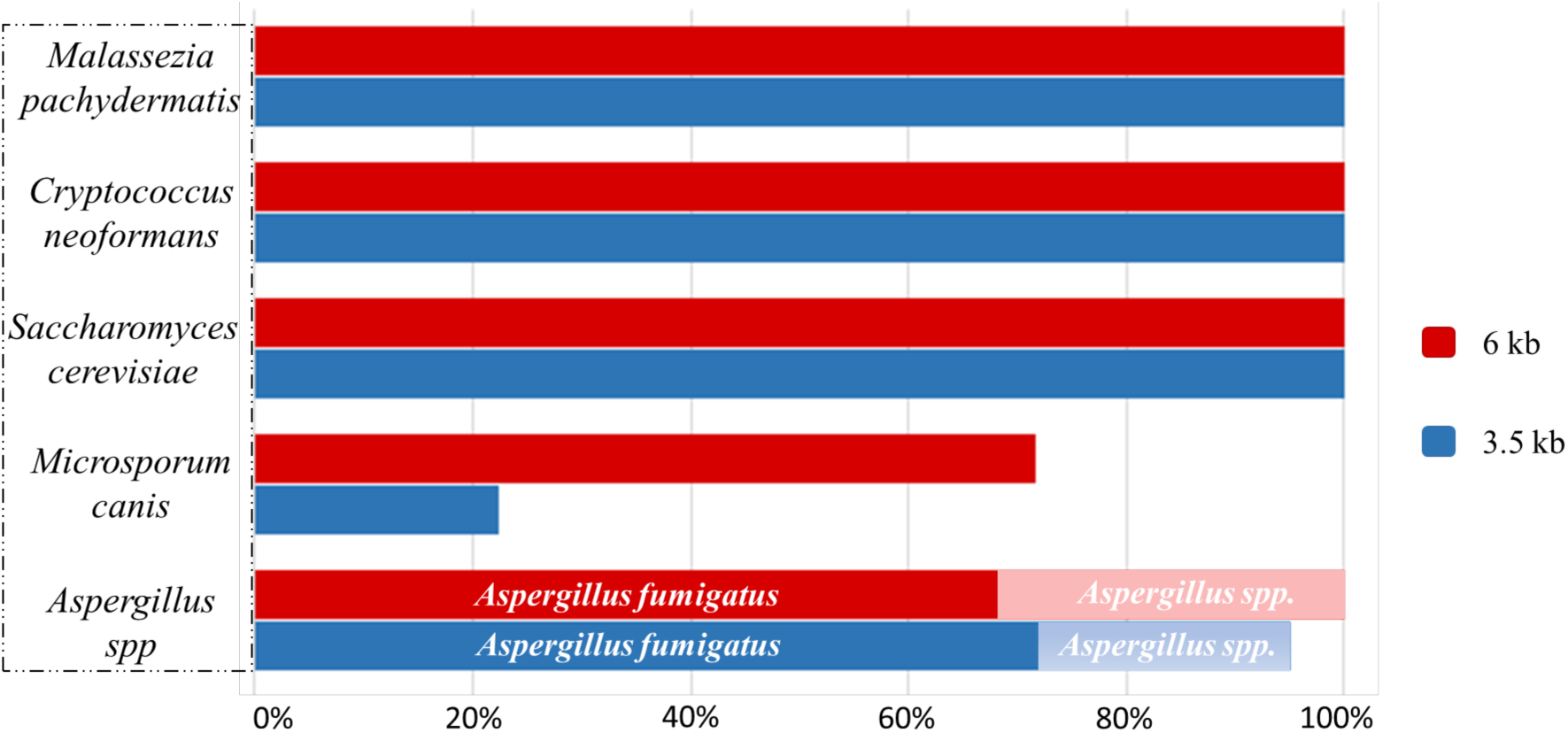
ZymoBIOMICS™ mock community (*S. cerevisiae* and *C. neoformans*) and microbiological cultures of fungi after taxonomical classification of the 3.5 Kb and 6 Kb ribosomal amplicons.

Both amplicons correctly detected and identified the ZymoBIOMICS™ mock community fungal strains (*Saccharomyces cerevisiae, Cryptococcus neoformans)*, and *Malassezia pachydermatis* and *Microsporum canis* from microbiological cultures. Looking in detail, *Saccharomyces cerevisiae, Cryptococcus neoformans* and *M. pachydermatis* were detected up to 100% by both 3.5 Kb and 6 Kb long fragments, while 6 Kb amplicon better detected *Microsporum canis.*

Both fragments identified *Aspergillus* genus as the main one found in the culture, but looking at the species level, *A. fumigatus* was the most abundant. *Alternaria* spp. and *Candida* spp. showed different results of what we expected (Figure 3).

**Figure 3.**
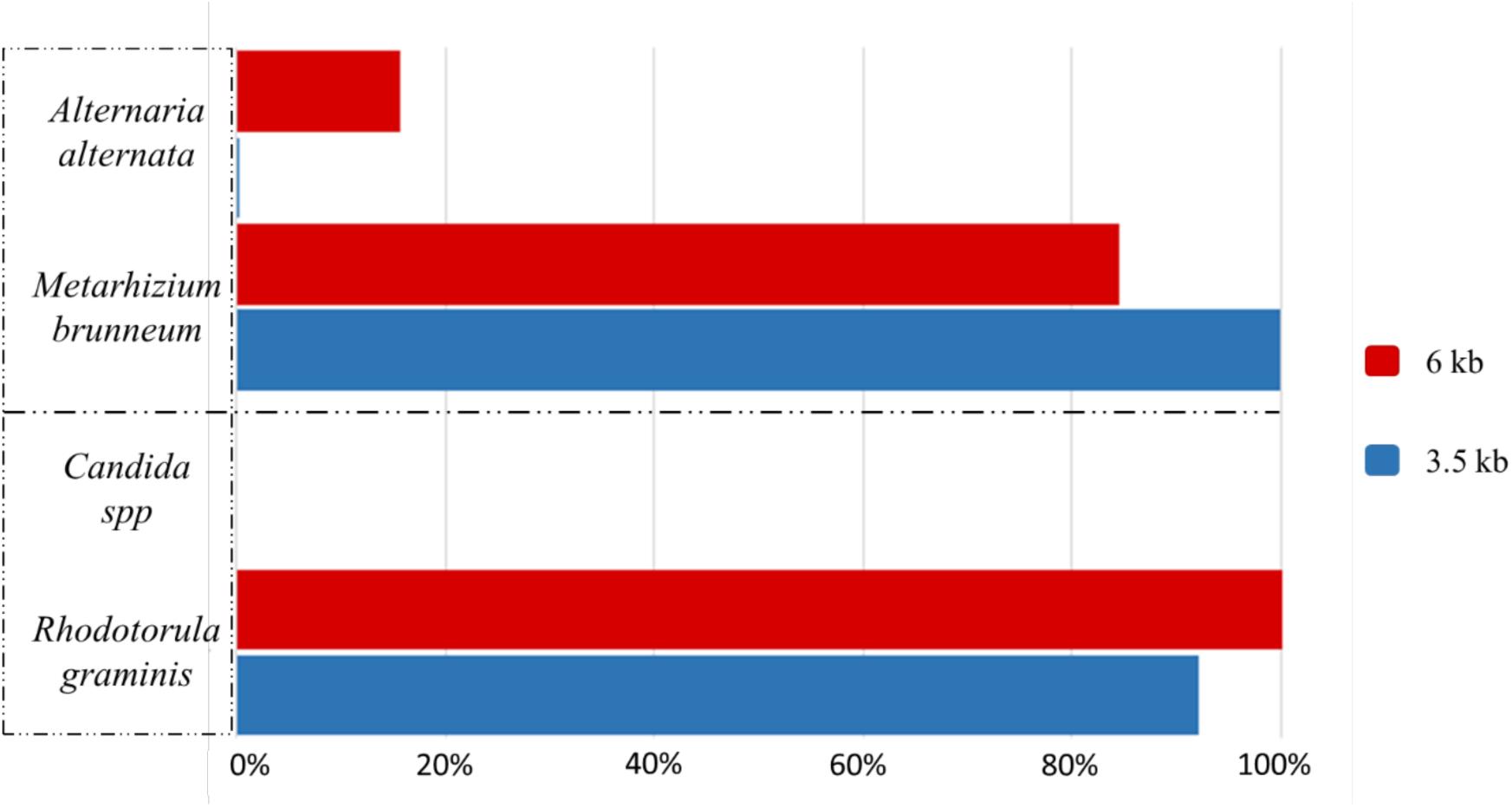
Fungal microbiological cultures showed unexpected results in the taxonomical classification after sequencing. Few reads from the *Alternaria* culture belonged to *Alternaria* spp, and it was classified at species level as *A. alternata*, but the most abundant fungus found was *Metarhizium brunneum*. No reads from the *Candida* culture were classified as *Candida* spp because of the presence of *Rhodotorula graminis*.

For the *Alternaria* culture, both amplicons size detected *Metarhizium brunneum* as the main fungus found in these cultures. Conversely, *Alternaria alternata* was the species found in really low relative abundance (0.2% for 3.5 Kb and 15.6% for 6 Kb). No similarities between these fungi were found: they belong to different order, Pleosporales and Hypocreales. *Alternaria* are ubiquitous filamentous fungi, which are found in soil air and human/animal skin (Pastor and Guarro 2008); *Metarhizium* is commonly found as a parasite of insects and symbiont of plants (Samish et al. 2014; Tiago and Oliveira 2014).

Looking at the nomenclature of this fungus, *Metarhizium brunneum* belonged to *Metarhizium anisopliae* strain (Tiago and Oliveira 2014; Yousef et al. 2018), but no correlation with *Alternaria alternata* was found.

For the *Candida* culture, the colonies of this fungus in Petri dish were red/orange. Sequences revealed the presence of *Rhodotorula graminis*, and only a few reads were classified as *Candida* spp. *Rhodotorula* is a carotenoid biosynthetic yeast, part of the Basidiomycota phylum, easily identifiable by distinctive yellow, orange or red colonies (Yadav et al. 2014). This yeast produces three major carotenoids: b-carotene, torulene and torularhodin, and is commonly associated with plants (http://www.antimicrobe.org/f16.asp#t1). In this case, we could confirm the presence of two yeast species in the microbiological culture.

### Canine Otitis

Conscious that no differences were found in *M. pachydermatis* analysis using both fragments sizes (Figure 2), we sequenced microbiological cultures of *M. pachydermatis* as positive controls and four complex samples with 3.5 Kb amplicon size. We run WIMP for fungal communities’ detection: the positive controls were identified as *M. pachydermatis*, while the complex otitis samples showed other *Malassezia* species (Table 2).

**Table 2:** Relative abundance of *Malassezia* species found in pure *M. pachydermatis* culture, and in four complex samples belonged to three different dogs affected by otitis. Samples S02 and S03 belong to the same dog, while S04 and S01 belong to two different dogs.

|  | *M. pachydermatis* | *M. globosa* | *M. sympodialis* | *M. Unidentified* | *Others* |
| --- | --- | --- | --- | --- | --- |
| M01 | 98.9% | 0.2% | 0.2% | 0.2% | 0.5% |
| M02 | 99.0% | 0.1% | 0.2% | 0.2% | 0.5% |
| M03 | 98.9% | 0.2% | 0.3% | 0.2% | 0.4% |
| S02_healthy | 98.2% | 0.1% | 0.2% | 0.2% | 1.3% |
| S03_affected | 95.1% | 1.1% | 1.7% | 0.4% | 1.7% |
| S04_affected | 78.8% | 4.0% | 5.0% | 3.9% | 8.3% |
| S01_affected | 73.7% | 4.7% | 5.1% | 5.3% | 11.2% |

Two of the samples correspond to the same dog, one from a healthy ear (S02) and the other one (S03) with clinical signs compatible with otitis externa, and *M. pachydermatis* is the main fungal species detected in both ears. The other two samples (S01 and S04) came from the ear with otitis externa of two dogs. In that case, other *Malassezia* species were detected together with *M. pachydermatis*, such as *M. globosa* and *M. sympodialis* (Table 2).

## DISCUSSION

Our first approach with Oxford Nanopore Technologies sequencing was aimed to understand if long amplicons are suitable markers to analyse the mycobiome in dog skin, and which size could be the best in the analysis of mycobiome. The microbiological cultures were essential for the study as positive controls because their genome sequences were used to validate the correct detection of fungi in WIMP.

Primers used to amplify the fungal ribosomal operon domains should be chosen depending on the fungus, but no standard markers are defined yet. The longest amplicons should be considered to describe the communities at lower taxonomy classification (Tedersoo et al. 2018; Wurzbacher et al. 2018). *Malassezia* spp, *Saccharomyces cerevisiae, Cryptococcus neoformans, Microsporum canis*, and *Aspergillus* spp, were correctly detected and identified from the microbiological cultures. However, the microbiological cultures corresponding to *Alternaria* spp and *Candida* spp were misidentified as per classical microbiology, and other fungi were detected. It is noteworthy that the samples plated came from dog skin, which is prone to environmental contamination, as has been previously described in skin microbiome of healthy dogs (Cuscó et al. 2017; Cuscó et al. 2019).

Few of the reads from the *Alternaria* culture were classified as *A. alternata* with the 6 Kb amplicon, while most classified as *Metarhizium brunneum*. Discovered in Spain and used as an herbicide against fly *Bactrocera oleae* (Yousef et al. 2018), this fungus belongs to the same phylum of Ascomycota, but it differs at lower taxonomy levels. The *Candida* microbiological culture was misclassified, even when showing an orange colour, caused by *Rhodotorula graminis*.

Finally, we investigate the possibility of reaching species level in complex samples from the skin of dogs affected by otitis, finding that *Malassezia* was the most abundant genus. The classification at the species level was performed to investigate possible changes between health status and diseased one.

*M. pachydermatis* has been reported as the most abundant species in the ear canal of healthy dogs (Korbelik et al. 2018). WIMP correctly identified all the *Malassezia* samples, and we were able to identify *Malassezia* at the species level from four complex canine otitis samples. Two of the samples corresponded to the same dog, one from a healthy ear (S02) and one with clinical signs (S03) that were compatible with otitis externa. *M. pachydermatis* is the main fungal species detected (98% of the reads for the healthy ear – S02 – and 95% for the sample with clinical signs compatible with otitis externa – S03). The other two samples (S01 and S04) came from the ear with otitis externa of two other dogs. In those cases, other *Malassezia* species were detected together with *M. pachydermatis*, such as *M. globosa* and *M. sympodialis*.

The results agree with previous studies on *Malassezia spp*, describing it as a commensal microorganism in human and animal skin that may become pathogenic (Cafarchia et al. 2005; Ngo et al. 2018).

In conclusion, this is a first approach to assess nanopore sequencing and the long-read amplicon approach for the analysis of fungi in complex samples. We assess the taxonomical power of two different amplicons targeting 3.5 Kb and 6 Kb of the ribosomal operon respectively, with the longest one providing a better taxonomy assignation. We used positive controls from the ZymoBIOMICS™ mock community, and from microbiological cultures that cannot always be considered pure cultures. We demonstrate the suitability of this approach to characterize the fungal community of complex samples, either healthy samples or samples with clinical signs of infection, such as otitis.

The next steps will lead to simplify the library preparation with the Rapid Barcoding kit from ONT and the analysis of complex samples from different origins to detect the causal agent of the disease in a clinical metagenomics approach.

## Supporting information

Dataset-FastQ

## Notes

https://zenodo.org/record/3662300#.XkUNZC3vXxp

### Competing Interest Statement

The authors have declared no competing interest.

