## Supplementary material for "Rapid and real-time identification of fungi up to the species level with long amplicon Nanopore sequencing from clinical samples": Dataset-FastQ

D'Andreano Sara, Cuscó Anna, & Francino Olga. (2020). Rapid and real-time identification of fungi up to the species level with long amplicon Nanopore sequencing from clinical samples [Data set]. Zenodo. http://doi.org/10.5281/zenodo.3662300
